# Actin force generation in vesicle formation: mechanistic insights from cryo-electron tomography

**DOI:** 10.1101/2021.06.28.450262

**Authors:** Daniel Serwas, Matthew Akamatsu, Amir Moayed, Karthik Vegesna, Ritvik Vasan, Jennifer M Hill, Johannes Schöneberg, Karen M Davies, Padmini Rangamani, David G Drubin

**Author notes:** Allen Institute of Cell Science, Seattle, WA, United States. Department of Pharmacology and Department of Chemistry & Biochemistry, University of California, San Diego, La Jolla, USA.

## Abstract

Actin assembly provides force for a multitude of cellular processes. Compared to actin assembly-based force production during cell migration, relatively little is understood about how actin assembly generates pulling forces for vesicle formation. Here, cryo-electron tomography revealed actin filament number, organization, and orientation during clathrin-mediated endocytosis in human cells, showing that force generation is robust despite variance in network organization. Actin dynamics simulations incorporating a measured branch angle indicate that sufficient force to drive membrane internalization is generated through polymerization, and that assembly is triggered from ∼4 founding “mother” filaments, consistent with tomography data. Hip1R actin filament anchoring points are present along the entire endocytic invagination, where simulations show that it is key to pulling force generation, and along the neck, where it targets filament growth and makes internalization more robust. Actin cytoskeleton organization described here allowed direct translation of structure to mechanism with broad implications for other actin-driven processes.

**Highlights:**

- Filament anchorage points are key to pulling force generation and efficiency.
- Native state description of CME-associated actin force-producing networks.
- Branched actin filament assembly is triggered from multiple mother filaments.
- Actin force production is robust despite considerable network variability.

## Introduction

Actin filaments are structurally polarized linear polymers that preferentially grow at one end, called the plus or barbed end (Pollard, 2016). Polymerization of individual actin filaments can produce forces in the range of 1 to 9 pN (Dmitrieff and Nedelec, 2016). These filaments can organize into higher order assemblies required for a multitude of essential cellular functions including clathrin-mediated endocytosis (CME) (Rottner et al., 2017). During CME, the plasma membrane is deformed to produce cargo-containing clathrin-coated vesicles (CCVs). This membrane remodeling is promoted by assembly of a clathrin-containing protein coat and forces provided by the actin cytoskeleton (Kaksonen and Roux, 2018). While how a growing actin network can push on a cellular membrane, for example during cell migration, is well understood, how assembly can aid in membrane pulling during endocytosis and intracellular trafficking is much less well understood. CME is well suited to studies of membrane pulling through actin assembly as nearly complete lists of the components involved and detailed information on their dynamics exist (Kaksonen and Roux, 2018). However, molecular-scale positional information about these components in their native state, which is essential for attaining a mechanistic understanding of their activities, is lacking.

During CME, clathrin coat assembly initiation is followed by recruitment of the actin filament nucleating Arp2/3 complex (Taylor et al., 2011). This complex can bind to the sides of existing “mother” filaments to induce assembly of new “daughter” filaments, leading to formation of branched actin networks (Rottner et al., 2017). The clathrin and plasma membrane-binding coat protein Hip1R can tether actin filaments to CME sites to harness filament polymerization forces for plasma membrane deformation (Akamatsu et al., 2020; Engqvist-Goldstein et al., 1999, 2001). Using agent-based models, we previously found that actin self-assembles into a branched network during CME, with filaments oriented orthogonal to the plasma membrane, surrounding, and attached to, the clathrin coat, and their growing plus ends oriented toward the plasma membrane. This geometry was dependent on the experimentally constrained spatial distribution of activated Arp2/3 complexes and actin filament-binding Hip1R linkers embedded in the clathrin coat (Akamatsu et al., 2020; Mund et al., 2018; Sochacki et al., 2017). The resulting network geometry was consistent with previously proposed models and predicted the range for the numbers of filaments that could be involved in force generation in CME (Akamatsu et al., 2020; Kaksonen et al., 2006). In contrast, platinum replica electron microscopy (EM) of unroofed mammalian cells showed branched actin filaments surrounding only the neck region of CME sites in a collar-like fashion with filaments aligned parallel to the plasma membrane (Collins et al., 2011). However, this method like other classic EM methods might not preserve native actin cytoskeleton organization as it might result in partial removal of some of the actin network during unroofing, and might not allow entire actin filaments in dense networks to be traced (Collins et al., 2011; Maupin-Szamier and Pollard, 1978; Resch et al., 2002; Small, 1981). Additional *in silico* experiments have shown a force requirement for actin polymerization in the range of <1 to 20 pN depending on the direction of force production and shape of the CME invagination, suggesting that both described types of filament arrangements could facilitate CME progression (Hassinger et al., 2017). These predictions emphasize the need to determine the orientation and numbers of actin filaments at CME sites before a quantitative understanding of the precise mechanism of actin-mediated force generation during CME can be achieved. The platinum replica EM study further suggested that the branched actin networks originate from a single founding mother filament, but its origin remains unclear (Collins et al., 2011). In addition to the orientation of branched actin filaments and the origin of mother filaments, the precise localization of the critical actin-CME linker Hip1R is ambiguous, since partially contradicting data has been published (Clarke and Royle, 2018; Engqvist-Goldstein et al., 2001; Sochacki et al., 2017). Obtaining the structural information described above requires a method that allows visualization of native 3D actin networks at the single filament level, determination of each filament’s orientation and the precise localization of the aforementioned linker protein Hip1R.

In recent years, *in situ* cryo-electron tomography (cryo-ET) has been shown to be an extremely powerful approach to visualize the three-dimensional organization of cellular features, including actin networks and protein-coated vesicles, at near native state conditions with unprecedented detail (Bykov et al., 2017; Fäßler et al., 2020; Jasnin and Crevenna, 2016; Mahamid et al., 2016; Vinzenz et al., 2012). Here, to elucidate the mechanism of pulling force generation through actin network assembly, we integrated cryo-ET on organization of actin networks involved in mammalian cell CME with mathematical modeling.

## Results

### Clathrin coat identification in cryo-electron tomograms of intact cells

To investigate the structural organization of actin during CME, we used cryo-ET of vitrified intact human-derived SK-MEL-2 cells growing on EM grids (Serwas and Davies, 2021). We identified honeycomb-like arrangements in our tomography data, which were reminiscent of clathrin coats seen in previous electron microscopy studies (Fig. 1A) (Avinoam et al., 2015; Cheng et al., 2007; Heuser, 1980). To test whether these arrangements are indeed clathrin coats, we applied correlative cryo-fluorescence light microscopy (cryo-FLM) and cryo-ET using SK-MEL-2 cells that endogenously express GFP-tagged clathrin light chain A (CLTA-GFP) and were incubated with fluorescent transferrin (TF) cargo to precisely pinpoint CME events (fig. S1, A and B). Tomograms obtained by this method showed the same structural features as in the randomly collected data sets (fig. S1B). The randomly collected tomograms were of higher quality than the correlative data, likely due to fewer manual handling steps, and were therefore used for the further analysis. Subtomogram averaging of clathrin-coat vertices from a total of 8 CME sites and clathrin-coated vesicles (CCVs) revealed additional details of the native hub structure (Fig. 1B and Movie S1). The resolution of our density map was 2.7 nm based on the 0.5 Fourier shell correlation criterion (Fig. 1C). Computational fitting of the recently published structural model obtained by single particle cryo-EM (PDB ID: 6SCT; (Morris et al., 2019)) into our *in situ* map resulted in a cross- correlation score of 0.94 (Fig. 1D). Our structural analysis together with our correlative microscopy results thus verifies the identity of the protein coats in our tomograms as clathrin coats.

**Fig. 1.**
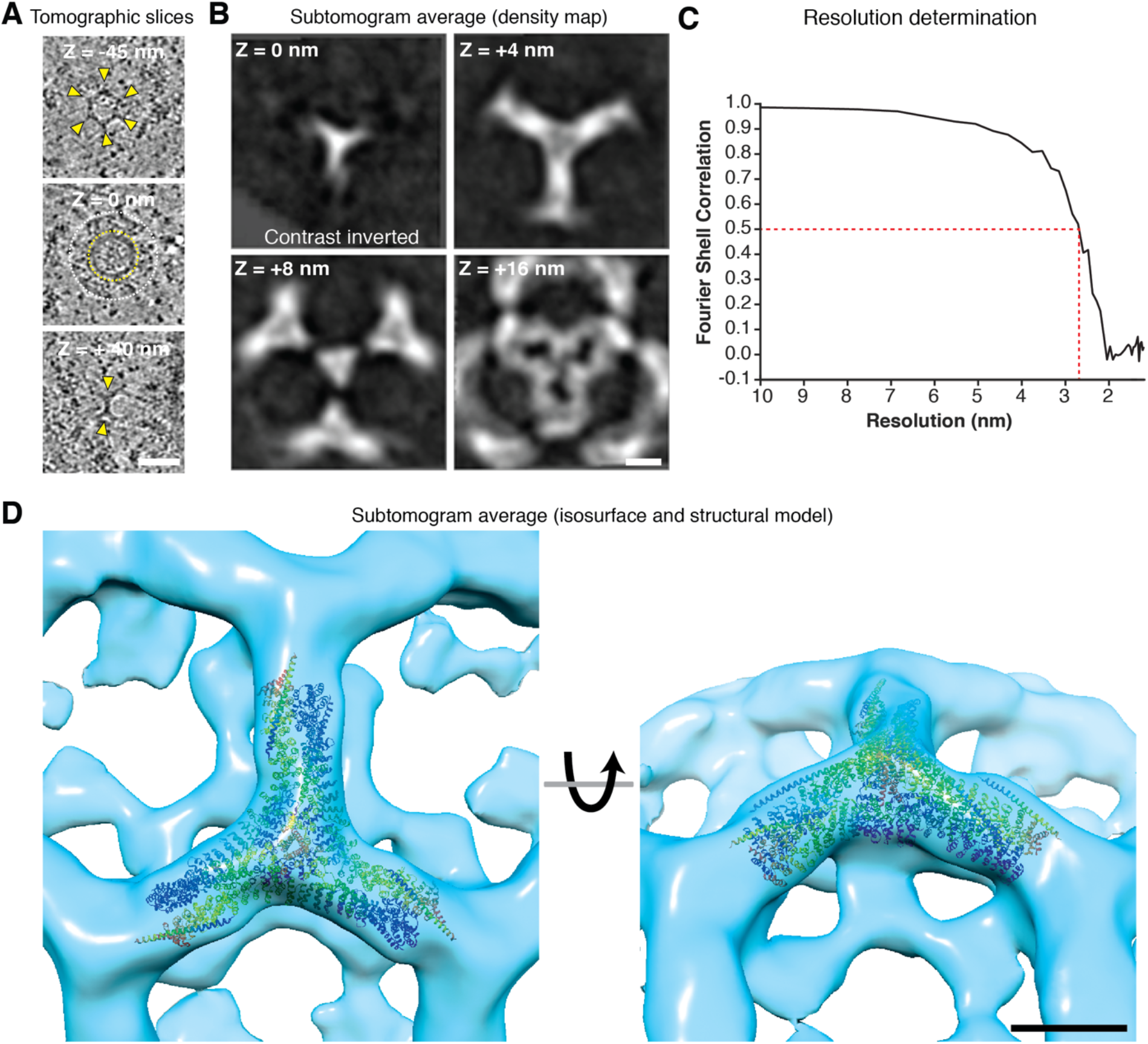
Native clathrin coat architecture and structure revealed by in situ cryo-ET. (**A**) Tomographic slices of a CCV at Z-positions relative to central slice (Z = 0 nm). Top and bottom slice show characteristic cage-like clathrin coat architecture with three leg-like extensions emanating from each vertex (yellow arrowheads). Yellow and white circles in central slice indicate vesicle and coated area respectively. (**B**) Density map of the clathrin coat determined within intact cells. (**C**) Resolution determination of the clathrin density map based on FSC. (**D**) Structural model of the clathrin hub (PDB ID: 6SCT, in rainbow colors, from (Morris et al., 2019) fitted into subtomogram average isosurface model of cytoplasmic clathrin vertex. Scale bars, 50 nm in (**A**), 10 nm in (**B**) and (**D**).

### Actin organization at different CME stages

During CME, the relatively flat plasma membrane invaginates and then a constricted neck forms at the base of the formed pit prior to CCV scission (Avinoam et al., 2015; Roth and Porter, 1964). We classified the clathrin structures in our tomograms according to the shape of their underlying membrane as early flat, early and late invaginated clathrin-coated pits (CCPs), and as CCVs (Fig. 2A, B, fig. S2A, B and Movies S2-S5). CCVs had a mean membrane and clathrin coat diameter of 46.7 ± 7.5 nm and 93.4 ± 4.2 nm respectively, while the CCP mean membrane and coat diameter were 90.2 ± 10.4 nm and 136.1 ± 17.7 nm (fig. S1C). We then generated segmentation models to visualize the spatial relationship between actin filaments, the membrane and the clathrin coat in 3D (Fig. 2B, and fig. S2B). Our analysis revealed that individual clathrin triskelia tended to be somewhat disconnected in early clathrin coats compared to late CME sites and CCVs, indicating flexibility of early clathrin coats, which is consistent with a recent study on unroofed, chemically fixed cells (Sochacki et al., 2021). Actin branch junctions could clearly be identified based on the presence of an arc-like density at the junctions, most likely representing the Arp2/3 complex branch nucleator (Fig 3A, and fig. S3A) (Rouiller et al., 2008; Vinzenz et al., 2012). To our surprise, CME-associated actin filaments did not consist of branched actin filaments exclusively as previously proposed (Collins et al., 2011). Instead, we observed a mixture of branched and unbranched filaments at all stages of CME (Fig. 2B, and fig. S2B). While branched filaments accumulated directly adjacent to CME sites, unbranched filaments were distributed across the entire tomographic volume. Unbranched filaments were organized in bundles close to CME sites, in dense meshworks of cortical actin filaments covering and surrounding CME sites, or as separated individual filaments. Branched actin filaments appeared asymmetrically distributed around CME sites, as suggested previously (Collins et al., 2011; Yarar et al., 2005).

**Fig. 2.**
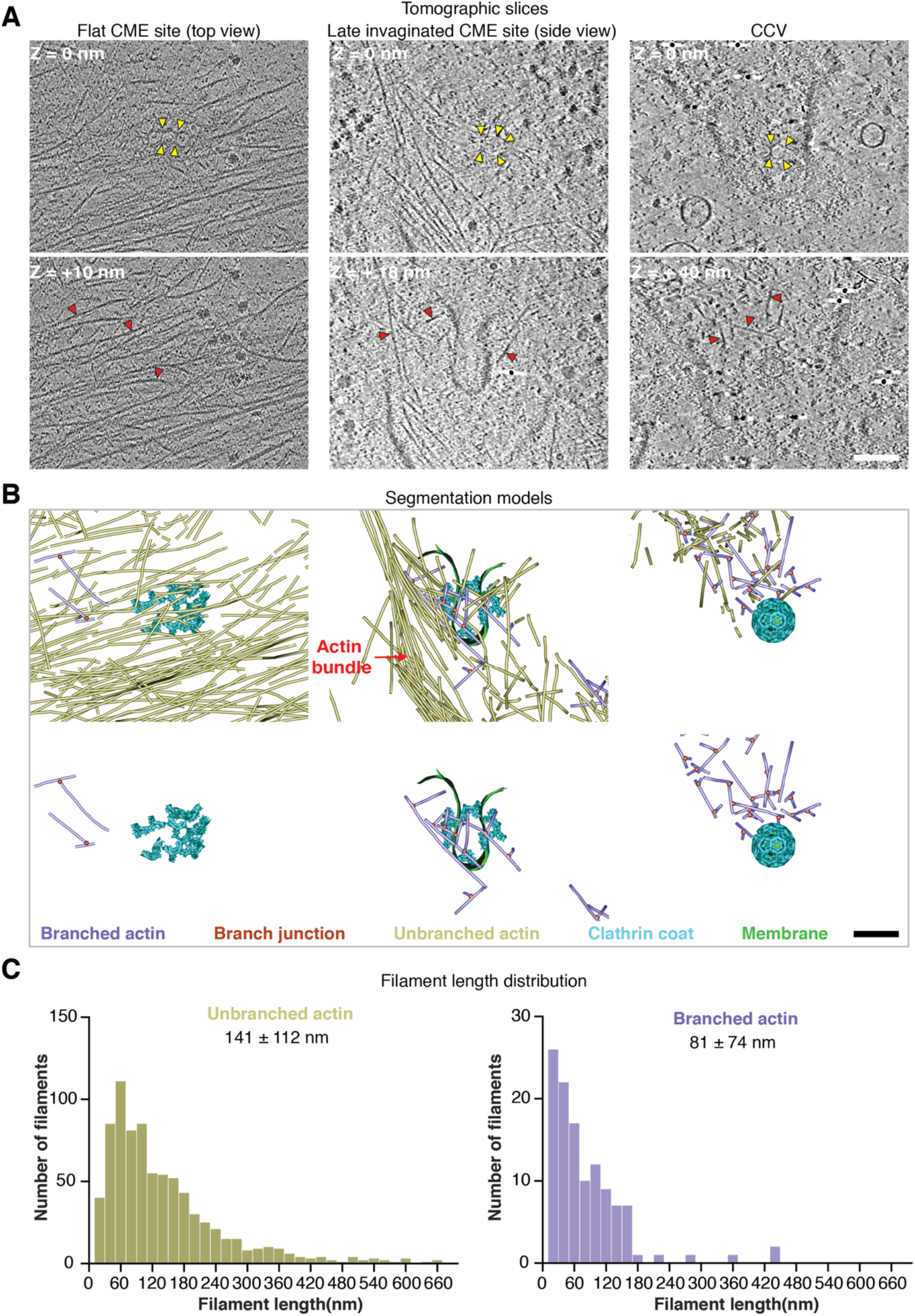
CME actin networks consist of branched and unbranched filaments. (**A**) Tomographic slices at Z-positions relative to central slice (Z = 0 nm) of CME events at indicated stages. Clathrin coat and individual actin filaments are highlighted with yellow and red arrowheads respectively. (**B**) Segmentation models of the tomograms in (**A**), bottom row shows branched actin filaments only. Color coded legend describes elements shown in the models. (**C**) Filament length distribution of unbranched and branched filaments across all CME events shown in this publication. Mean filament length and SD for unbranched and branched actin filaments are highlighted. Scale bars, 50 nm in (A, B).

**Fig. 3.**
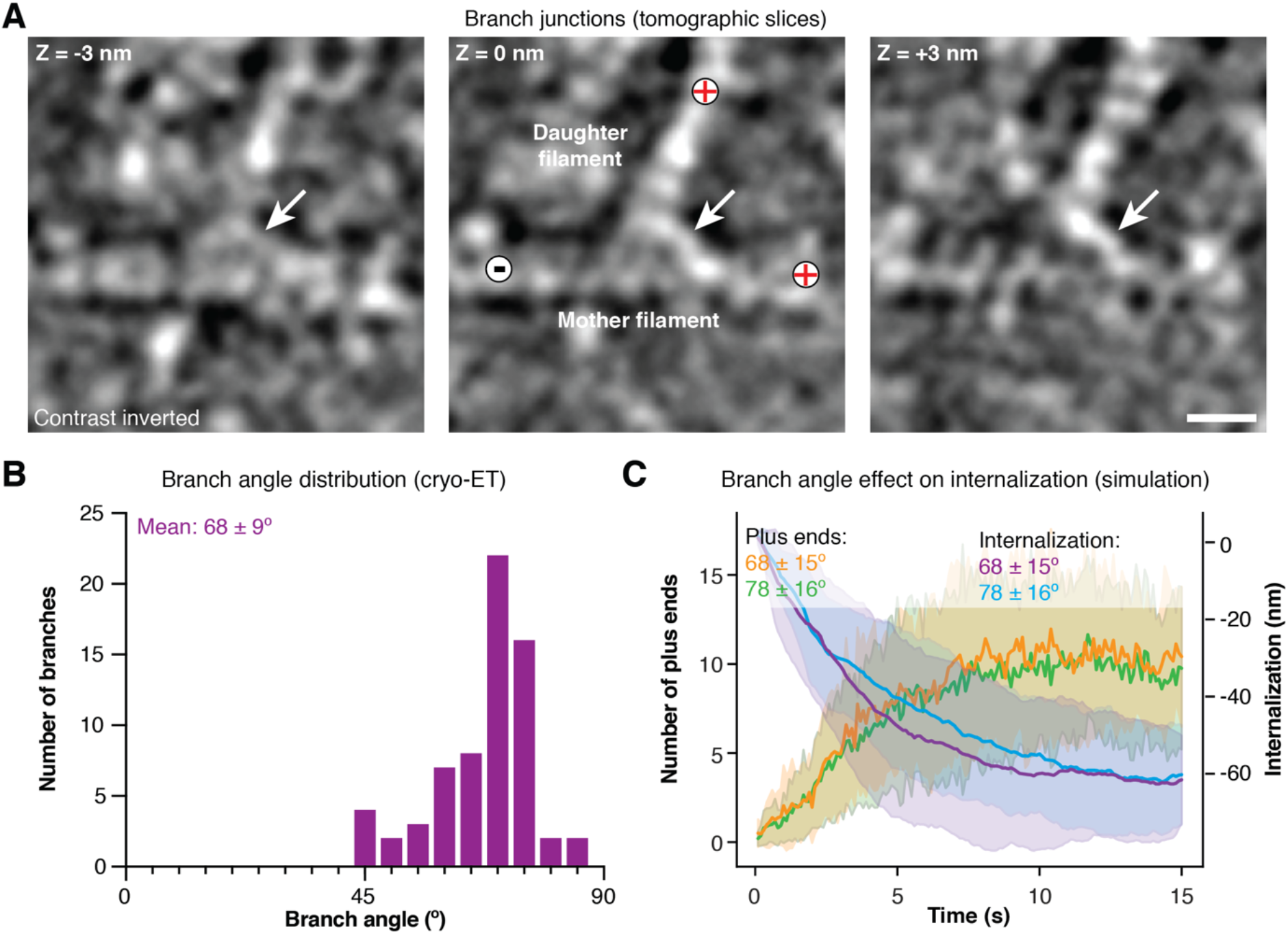
CME is robust against branch angle variation. (**A**) Tomographic slices of an individual branch junction at Z-positions, left to right, relative to central slice (Z = 0 nm). Positions of mother, daughter filaments, plus and minus ends are indicated. Arrow points to the arc-like density of the Arp2/3 complex. (**B**) Branch angle distribution measured in tomograms. Mean value and standard deviation are displayed. (**C**) Result of simulations showing the effect of branch angle variation on the number of plus ends at the base of CME invaginations (green and orange curves) and the internalization rate (cyan and purple curves). Mean and SD are shown. Scale bar, 5 nm in (**A**).

The mechanical properties of individual actin filaments depend on their length, with shorter filaments being more rigid than longer ones (De La Cruz and Gardel, 2015). Our previous simulations predicted that branched filaments grow to a length of 90 ± 80 nm during CME (Akamatsu et al., 2020). Here we measured an average branched filament length in tomograms of 81.74 ± 74 nm, indicating that these filaments could have assembled over the course of a CME event (Fig. 2C). This possibility is further supported by the relatively low number of branched filaments in early-stage CME tomograms shown in Fig. 2A, B. Unbranched filaments were on average longer (141 ± 112 nm), which suggests that some filaments might have existed before the onset of the CME event. However, many unbranched filaments fall in the same length category as the branched filaments, which indicates that these could have been newly polymerized and also contributed to force generation to support CME progression (Fig. 2C).

### CME is robust to variations in actin filament branch angle

A wide range of branch angles, obtained from various sample types and methods, have been reported previously for Arp2/3 complex-nucleated actin filament networks (Blanchoin et al., 2000; Fäßler et al., 2020; Jasnin et al., 2019; Mueller et al., 2014; Mullins et al., 1998; Rouiller et al., 2008; Svitkina and Borisy, 1999; Vinzenz et al., 2012). Here we measured an average branch angle of 68 ± 9°, which is in close agreement with recently published *in situ* structures (Fig. 3B) (Fäßler et al., 2020; Jasnin et al., 2019). We could not find a strong effect of the “missing wedge” characteristic of tomography data on our measurements (fig. S3B, see Material and Methods for details) (Wan and Briggs, 2016). The native branch angle was smaller and the junctions slightly stiffer (indicated by the standard deviation) compared to the average branch angle of 78 ± 16° in our previous mathematical model (Akamatsu et al., 2020). To test whether the smaller branch angle affects the force production capability, we conducted simulations of actin assembly at CME sites using the newly measured branch angle (fig. S3C). We found that for both ranges of branch angles, the number of plus ends polymerizing against the base of the CME site remained the same (10 ± 4) leading to similar internalization rates (Fig. 3C). We conclude that polymerization-based force production for CME is robust against branch angle variation.

### Branched actin assembly is nucleated from multiple mother filaments

Next, we tested the previously postulated notion that branched actin network assembly at CME sites is initiated from a single “founding” mother filament (Collins et al., 2011). Our model predicted that branched actin network assembly is initiated from several “founding” mother filaments, each giving rise to a distinct branched actin cluster. The number of clusters and number of filaments per cluster increased during CME progression to an average of 4 ± 2 clusters with 49 ± 21 filaments in each cluster (Fig. 4A, B and fig. S4A). The predicted clustered branched actin filament organization is consistent with the arrangement of branched actin filaments in the tomograms, where the average cluster number was 8 ± 6 (Fig. 3C, D). However, individual clusters only consisted of an average of 2.2 ± 0.5 filaments in the tomography data (fig. S4B). The discrepancy in the filament number per cluster in the model vs experiment can be understood as follows: even though the model shows 49 ± 21 filaments per cluster at the end of a CME event, the number of filaments at the base is only 10 ± 4, suggesting that not all filaments in the model contribute directly to force generation (Fig. 3C). Since the model only captures select actin interactions, it is also possible that the excess filament number per cluster in the model reflects certain limitations in the model. For example, in the current version of the model, the spacing between individual branches along a filament cannot be controlled. This might have led to branching events that may not be feasible in cells due to geometrical and cytoplasmic crowding-related spatial restrictions. The early CME site in the tomogram in Fig. 2 showed the lowest number of branched actin filament clusters, suggesting that more branched actin clusters are initiated and assembled during the plasma membrane internalization phase.

**Fig. 4.**
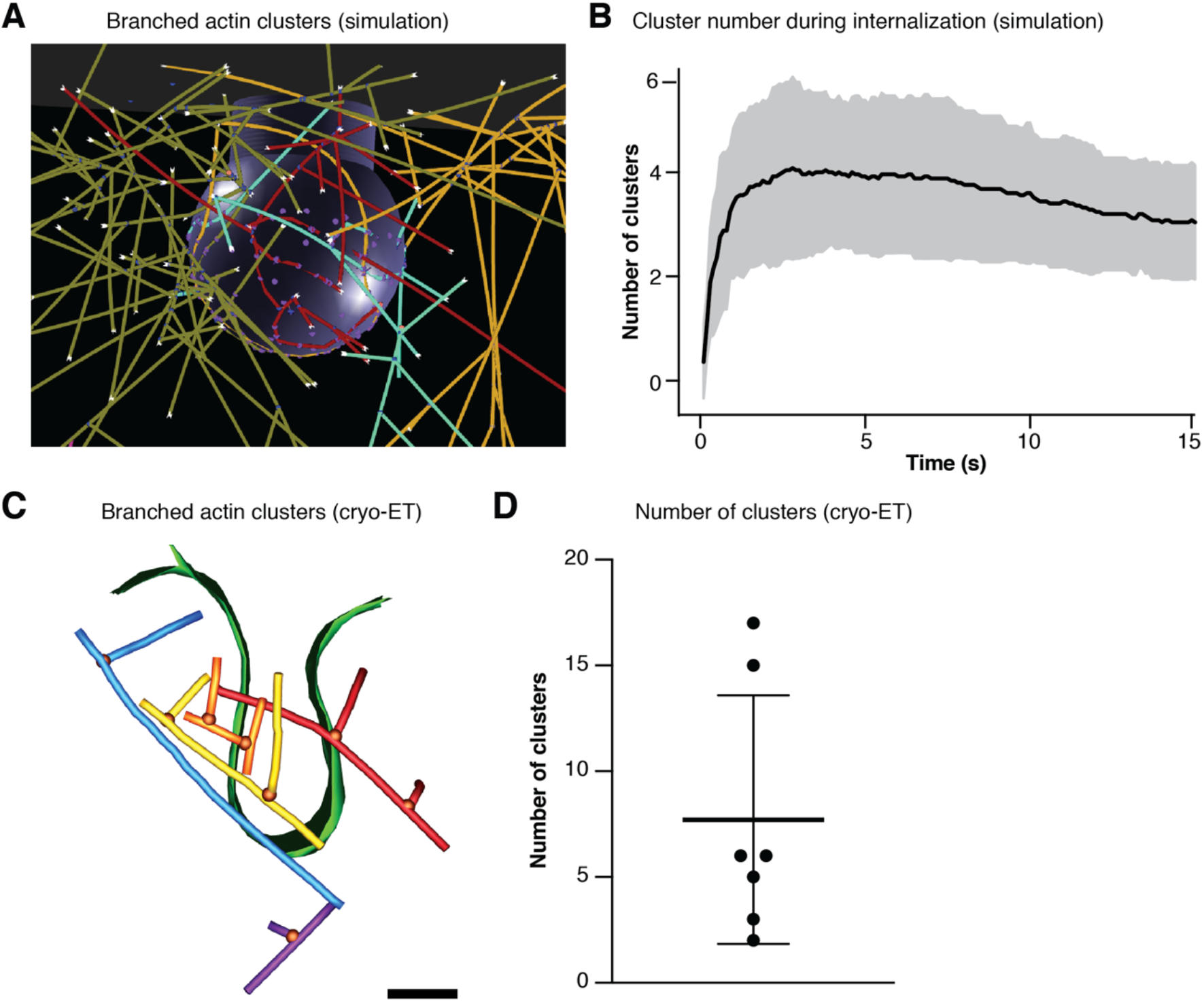
Branched actin filaments are organized into clusters. (**A**) Simulation snapshot highlighting clustered organization of branched actin filaments in mathematical model. Individual clusters are color coded. (**B**) Simulation of mean cluster number during internalization. (**C**) Segmentation model of late CME site from Fig. 2A highlighting clustered organization of branched actin filaments. Individual clusters are color coded. (**D**) Number of clusters found at individual CME events. Mean and SD are shown. Scale bar 50 nm in (**C**).

### Actin filament orientation at CME sites

Next, we set out to analyze the polarity and orientation of the actin filaments in the tomograms to assess the direction of force production through polymerization. The polarity of branched filaments can be determined based on branch junction geometry (Fig. 3A) (Narita et al., 2012). To analyze the polarity of unbranched filaments, we adapted a previously published method, which is based on cross-correlation analysis of the filaments in tomograms against simulated reference filaments with known polarity (fig. S5A-E) (Narita et al., 2012). We developed an analysis software package that allowed us to calculate and plot the orientation of the filaments relative to the normal vector of a simulated reference plane representing the position of the plasma membrane or individual CCVs. Orientation of the vector was defined such that 0° orientation indicates actin filament plus end pointing toward the reference plane and 180° orientation indicates actin filament plus end pointing away from the plane. The average filament orientation across all tomograms was 86 ± 31° for unbranched and 81 ± 36° for branched filaments, indicating that a large proportion of filaments were oriented parallel to the plasma membrane (fig. S5F, G). Further examination of individual CME sites revealed two general orientation types for branched actin filaments in close proximity to membrane invaginations, one in which filaments were oriented orthogonal to the plasma membrane, similar to the predictions from our simulations, and a second in which filaments were oriented largely parallel to the plasma membrane, similar to what was observed in previous work using platinum replica EM (Fig. 5A-D) (Akamatsu et al., 2020; Collins et al., 2011). On average 11 ± 8 branched and 36 ± 7 unbranched plus ends were pointing toward the plasma membrane (relative angle < 90°) during the membrane internalization phase for both types of filament arrangements (Fig. 5E). In contrast to our simulations, we also observed filaments oriented with their plus ends toward the neck of CME invaginations for both types of actin arrangements, where they could potentially produce a squeezing force to support scission (Fig. 5B and C). We also found branched actin filaments oriented with their plus ends towards CCVs, suggesting a role for filament polymerization in CCV transportation inside cells (fig. S5H, I).

**Fig. 5.**
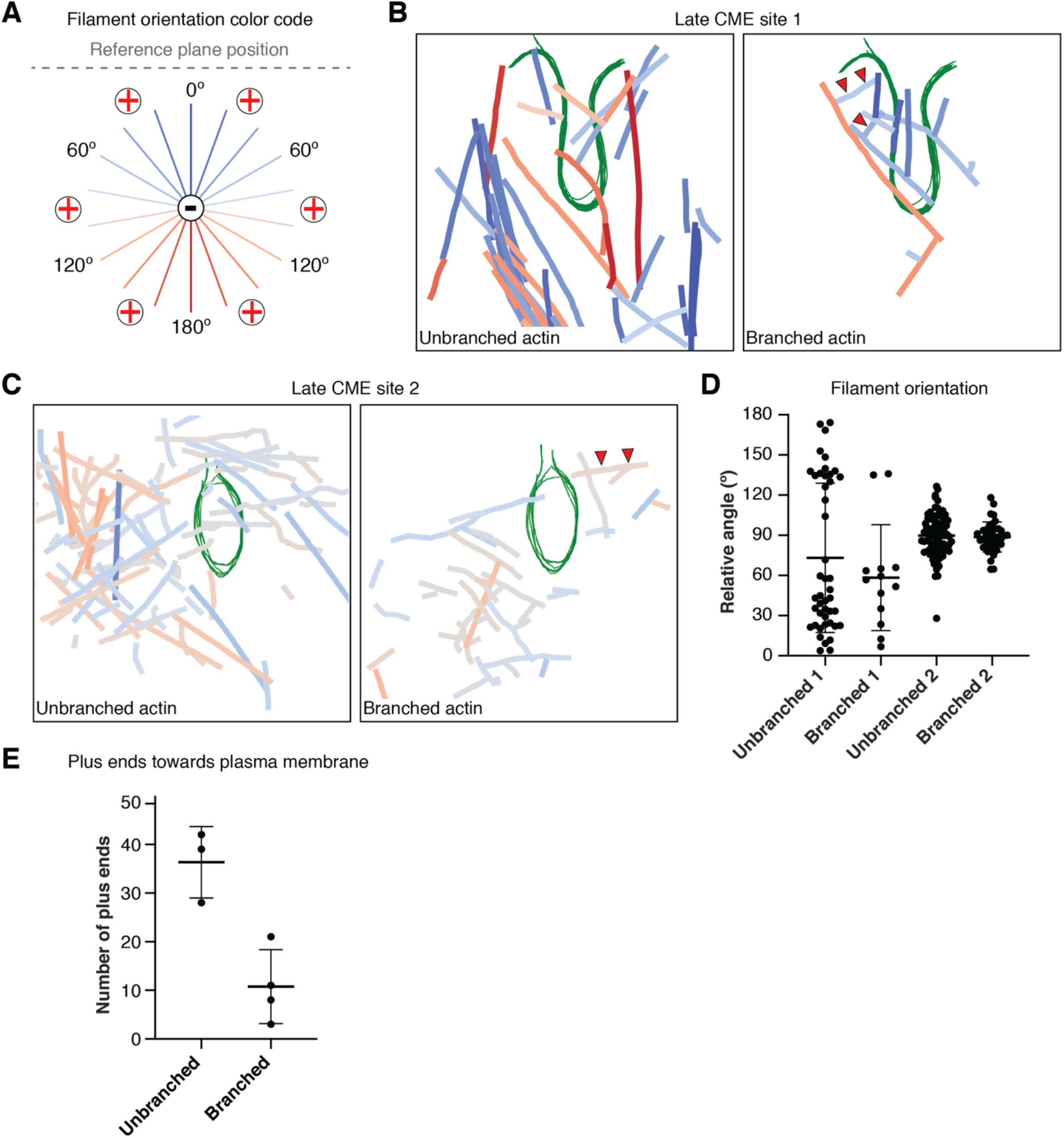
Actin filament orientation at CME sites. (**A**) Color code indicating filament orientation relative to the normal (0°) of a reference plane representing the plasma membrane. At 0°, filament plus ends point towards the membrane. (**B**, **C**) Color-coded filament orientation at two late-stage CME invaginations (green), (**C**) shows a site towards the end of vesicle scission. Note the presence of filaments with their plus ends oriented towards the neck region (red arrowheads). (**D**) Relative filament to reference plane orientation of late CME site 1 and 2 shown in (**B**) and (**C**), respectively. (**E**) Number of filament plus ends pointing towards the plasma membrane. Mean and SD are shown in (**D**) and (**E**).

### Hip1R localizes to the CME neck where it is predicted to increase internalization efficiency

Our previous model identified the localization and distribution of the actin binding protein Hip1R as a main determinant for actin organization and function (Akamatsu et al., 2020). There, we had assumed that Hip1R exclusively localizes to the tip of CME invaginations, based on (Sochacki et al., 2017), in which averaging of fluorescence light microscopy data was used (Akamatsu et al., 2020; Sochacki et al., 2017). However, individual images from (Sochacki et al., 2017) and other published work show variable Hip1R localization, including at the neck of CME invaginations (Clarke and Royle, 2018; Engqvist-Goldstein et al., 2001; Sochacki et al., 2017). Importantly, only the methodology in (Clarke and Royle, 2018) allowed visualization of the entire CME invagination and it is likely that the other localization data gave an incomplete picture. Therefore, we asked if we could identify Hip1R in our tomograms and whether its localization might play a role in targeting actin filaments to the neck of CME sites. Hip1R forms dimers that by quick-freeze deep-etch EM appear as ∼60 nm rod-shaped densities with two globular actin-binding domains at one end and two globular membrane-binding domains at the other (Engqvist-Goldstein et al., 2001). We identified densities resembling Hip1R in size and structure over the invagination surface including the neck of CME invaginations as well as at the plasma membrane adjacent to the pit, which was consistent with work from (Clarke and Royle, 2018) (Fig. 6A and fig. S6A). To further test whether these densities could be Hip1R dimers, we subjected the putative cytoplasmic actin binding domains to subtomogram averaging, yielding a low-resolution density map (Fig. 6B and fig. S6B, C). The size and shape of the density was sufficient to house two actin-binding domains plus the dimerization domains of the homologous protein talin and to fit onto an actin filament (Fig. 6B) (Gingras et al., 2008). This analysis provides further support for the likelihood that these densities are Hip1R dimers. We then tested in our model the consequence of Hip1R localization at the neck by conducting simulations wherein Hip1R molecules were positioned along the neck surface in addition to the invagination tip. The number of filament plus ends at the base (10 ± 4) remained the same with or without additional Hip1R dimers at the neck. However, additional 4.5 ± 2 plus ends were now found in the neck region (Fig. 6C and fig. S6D, E). Strikingly, in our simulations Hip1R neck localization not only directed filaments to the neck region, but it also strongly improved internalization efficiency, indicating that Hip1R at the neck can help with actin-mediated force generation (Fig. 6D).

**Fig. 6.**
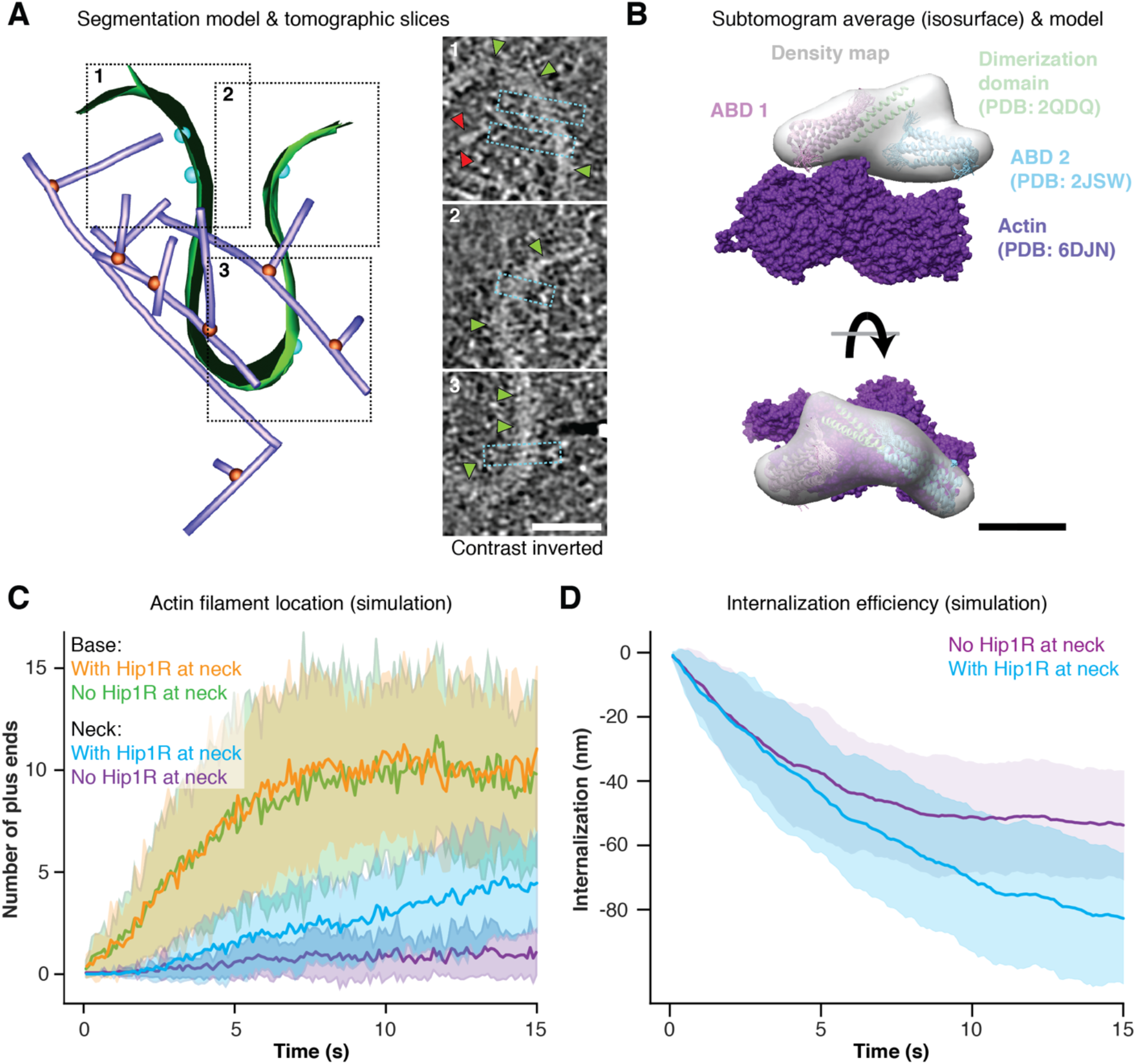
Hip1R-like linkers observed at the neck of CME invaginations increase internalization efficiency. (**A**) Tomographic slices of the regions indicated in the segmentation model of the tomogram shown in Fig. 2 (**A**), (**B**). Slices highlight putative Hip1R dimers (cyan boxes) that are associated with the coated membrane of the CME invagination (green arrowheads) and their positions are highlighted by cyan spheres in the segmentation model correspond. Red arrowheads in slice 1 point at an actin filament associated with a putative Hip1R dimer. (**B**) Superimposition of the surface model of the density map in (fig. S6 B) with previously published atomic models of two actin binding domains (ABD; PDB ID: 2JSW, (Gingras et al., 2008)) and two dimerization domains (PDB ID: 2QDQ, (Gingras et al., 2008)) of the homologous protein talin. Model of actin filament (PDB ID: 6DJN, (Chou and Pollard, 2019)) was added as a reference. Mathematical model readout showing filament distribution between CME invagination base and neck in the presence or absence of Hip1R molecules at the neck. (**C**) Result of simulations showing the effect of Hip1R neck localization on the number of plus ends at the base (orange and green curves) and neck region (cyan and purple curves) of CME invaginations. Mean and SD are shown. (**D**) Simulated internalization efficiency with and without Hip1R molecules at the neck. Scale bar, 50 nm in (**A**) and 5 nm in (**B**).

## Discussion

Forces produced by actin filament assembly are harnessed for many membrane remodeling processes, but much remains to be learned about the underlying assembly regulation and force- producing mechanisms. CME is a particularly attractive process for elucidating these mechanisms because it consists of a predictable series of morphological changes, each of which is coupled to recruitment of specific proteins. This work set out to define the three-dimensional organization of individual actin filaments at different CME stages because this information directly informs the force producing mechanism. What follows is an attempt to synthesize our cryo-ET and simulation results into a harmonious model for how actin assembly forces are generated and harnessed during CME.

Our finding that branched actin networks are organized in branched actin filament clusters at individual CME sites supports the conclusion that these clusters originate from multiple founding mother filaments (4 ± 2 in the simulations and 8 ± 6 in the tomograms), which is a different conclusion from the single mother filament model that was suggested previously (Collins et al., 2011). This clustered organization may provide flexibility for branched actin filaments to assemble in a crowded environment like the cell cortex, which also suggests that excluded volume effects may affect branched actin network assembly (Schreiber et al., 2010). The number of filaments and branched actin clusters was greater during the internalization phase and branched actin filament length was longer, indicating that these filaments polymerized over the course of a CME event and are thus capable of providing assembly force (Akamatsu et al., 2020). Variation in branched actin filament organization and in the number of clusters and filaments between individual CME sites likely reflects the stochastic nature of filament assembly as well as local adaptive responses to variable conditions. High variability in actin network organization and density from one CME site to the next might result from such factors as variance in mother filament number and orientation, Hip1R localization, active Arp2/3, N-WASP position and differences in plasma membrane tension (this study, (Akamatsu et al., 2020; Clarke and Royle, 2018; Engqvist-Goldstein et al., 2001; Kaplan et al., 2021; Sochacki et al., 2017)).

Work in the CME field has mainly focused on assembly of branched actin networks and did not take the presence of unbranched filaments at CME sites into account when investigating actin function. Also, unbranched actin filaments were not identified by the previous EM work, likely due to the inability to trace every individual filament along its entire length by platinum replica EM (Collins et al., 2011). Given the high number of unbranched actin filaments identified in our tomograms, two important questions are: Where do they come from and what are their functions? The presence of unbranched filaments at early stages of CME suggests that they represent pre- existing cortical actin filaments (early-stage site in Fig. 2, CME invaginations in fig. S2) or filaments originating from other close-by actin structures like the bundle in the late-stage tomogram in Fig. 2. These dense cortical actin arrangements might represent a physical barrier that needs to be cleared by filament severing, disassembly or repositioning for CME progression. Shorter filaments might be ones that are diffusing through the cytoplasm (Chen and Pollard, 2013; Raz-Ben Aroush et al., 2017). Importantly, some of the unbranched filaments might be actively producing assembly forces at CME sites. This possibility is supported by the presence of the Arp2/3 nucleation promoting factor SPIN90 at CME sites, which promotes nucleation of unbranched actin filaments and plays a role in epidermal growth factor receptor endocytosis (Kim et al., 2006; Luan et al., 2018; Oh et al., 2013). In all of these scenarios, unbranched actin filaments represent a potential resource for mother filaments required for Arp2/3-mediated, SPIN90- independent, branched actin filament nucleation (Pollard, 2007). The abundance of multiple sources of available mother filaments and pathways for actin filament assembly may act as a safety cushion that allows actin adaptation to ensure robust CME under variable conditions within a cell.

Because our cryo-ET approach allowed individual actin filaments to be fully traced and oriented in the volume surrounding CME sites, we were able to use simulations to assess the force-producing capabilities of these networks. We found that on average during invagination formation and neck constriction stages, 11 ± 8 branched and 36 ± 7 unbranched filaments were oriented with their plus ends toward the plasma membrane (relative angle < 90°), and additional filament plus ends were oriented toward the neck region (Fig. 5). In the simulations, we found that 10 ± 4 filament plus ends assembled at the base of CME invaginations, which was sufficient for successful internalization (Fig. 3C and 6C, D; (Akamatsu et al., 2020)). In previous simulations, the number of plus ends varied between 2 at low and 22 at high membrane tension conditions (Akamatsu et al., 2020). The force requirement to support the transition from a U-shaped invagination to an omega shape with a constricted neck through actin polymerization was predicted to be < 1 pN when the force is applied in parallel to the plasma membrane and 15 pN when applied orthogonally (Hassinger et al., 2017). Polymerization of individual actin filaments can provide between 1 and 9 pN of force (Dmitrieff and Nedelec, 2016). Despite the variability in actin network organization observed in the tomograms, our simulations predicted that the internalization efficiency was robust under different conditions. This conclusion suggests that actin filament assembly can harness the local variability and stochasticity at the filament level to generate efficient, robust force-generating machineries at the network level for CME. The actin-binding linker Hip1R might provide a spatial constraint for actin assembly to ensure robust internalization. Here, we find that Hip1R is not only at the tip surface of the CME invagination, but also at the neck region. In simulations, Hip1R neck localization directs filaments toward the neck constriction and improves internalization efficiency. Actin filament growth towards the neck also supports a role for actin polymerization during vesicle scission.

The principles identified here are expected to apply to other actin driven processes where cellular membranes are being pushed, pulled or squeezed (Fig. 7). The finding that branched actin filament networks at individual CME sites are organized in multiple discrete clusters is similar to force producing branched actin assemblies in lamellipodia, which push the plasma membrane outwards during cell migration (Vinzenz et al., 2012). However, presence of actin filament anchoring points at CME sites allows conversion of pushing force into pulling and squeezing forces. Internalization efficiency and filament orientation strongly depend on the distribution of these anchor points (this study and (Akamatsu et al., 2020)). The same mechanism might facilitate budding and fission at intracellular membranes, for example during vesicle budding from the trans-Golgi, where Hip1R anchoring points are important, or endosomes where the actin- and lipid-binding protein moesin could mediate anchorage (Carreno et al., 2004; Fehon et al., 2010; Muriel et al., 2016). Actin filaments in filopodia are also anchored to the plasma membrane, which might be important for filopodia formation and maintenance (Medalia et al., 2007). However, instead of being pulled inwards, the plasma membrane is pushed outwards. Besides of the position of anchoring points, we identified the position of actin filament assembly factors as a second constraint that defines the geometry of CME-actin networks, which is also likely to be the case for filopodia extension (Akamatsu et al., 2020). The nucleation promoting factor N-WASP arranges in a ring around CME sites (Almeida-Souza et al., 2018; Mund et al., 2018). In contrast, actin assembly factors accumulate at the tip of filopodia (Rottner et al., 2017). Interestingly, previous structural work also suggests that, similar to our results, about 10 filaments are involved in filopodia extension of (Medalia et al., 2007). Moreover, analysis of baculovirus-induced actin comet tails by electron tomography of membrane extracted cells suggests that the intracellular pathogen is pushed through the cytoplasm by the simultaneous assembly of 2 – 6 branched actin filaments (Mueller et al., 2014). We found that endocytic vesicles are pushed through the cytoplasm by a comparable number of branched actin filaments (fig. S5H, I).

**Fig. 7.**
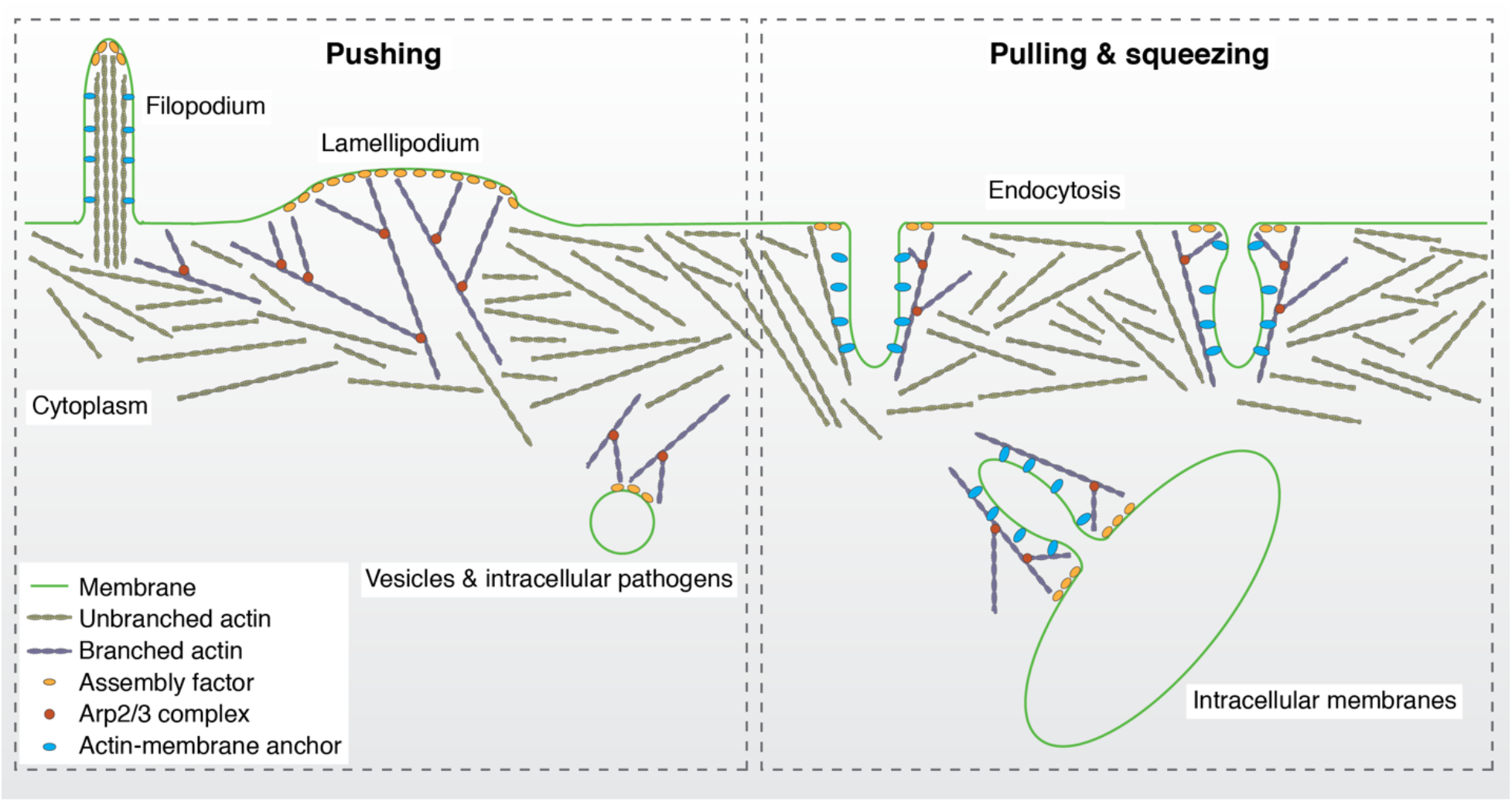
Harnessing actin polymerization force for the pushing, pulling and squeezing of cellular membranes. Actin polymerization can generate a pushing force to extend the plasma membrane during the formation of filopodia or lamellipodia, and to push vesicles and intracellular pathogens through the cytoplasm. Proteins with both actin- and membrane-binding abilities (e. g., Hip1R) can anchor actin filaments to the membrane, converting pushing force into pulling and squeezing forces based on their spatial localization. In addition to the distribution of anchoring points, actin network morphology and polymerization force direction are defined by a second geometrical constraint, the position of actin assembly factors. Note that the precise localization of assembly factors and actin-membrane linkers is not clear for most of the displayed processes.

In conclusion, we provided a comprehensive description of the complex native actin architecture during mammalian CME in unprecedented detail. We have precisely quantified the filament number and their orientation at CME events allowing for unambiguous predictions on their force production capabilities, which has not been achieved for other actin-driven processes in unperturbed cells. Our experimental and modeling data both highlight the remarkable flexibility and robustness of productive actin network organization. We conclude that assembly of actin networks in human cells produces sufficient force to account for robust internalization and neck constriction. By combining structural analysis and mathematical modeling, we gained previously inaccessible mechanistic insights into CME actin regulation and function, which has implications for many other actin-driven processes.

## Materials and Methods

### Cell culture

Double tagged SK-MEL-2 cells endogenously expressing clathrin light chain CLTA-TagRFP and dynamin2-eGFP, and single tagged SK-MEL-2 cells endogenously expressing clathrin light chain CLTA-eGFP, were cultured in DMEM/F12 (Gibco™) supplemented with 10% fetal bovine serum (premium grade, VWR Life Science Seradigm) and 1% Penicillin/Streptomycin (Gibco™). The double-tagged cell line was generated using zinc-finger nuclease-mediated genome editing and was published previously (Doyon et al., 2011; Grassart et al., 2014). The same reagents were used for the single-tagged cell line, except that the TagRFP in the donor plasmid used to construct the double tagged cell line was replaced with eGFP (Doyon et al., 2011; Grassart et al., 2014). Genome-edited cells were authenticated by short tandem repeat profiling. SK-MEL-2 cell lines were used because of their robust endocytic dynamics, good adherence and spreading behavior on EM grids.

### Cryo-sample preparation

Holey carbon grids (Quantifoil R2/1, 200 mesh, gold) were washed in acetone while agitating on a rocker for 15 to 30 min to remove potential residual plastic backing, washed in H_2_O and dried. Grids were then glow discharged using a Pelco SC-6 sputter coater, sterilized in 70% ethanol in H_2_O. Grids were placed in a 6-well plate containing cell culture medium (one grid per well) under sterile conditions in a biosafety cabinet. Grids were incubated in cell culture medium at 37°C and 5% CO_2_ in a cell culture incubator overnight. Cell culture medium was replaced the next day. SK-MEL-2 cells from a 75–90% confluent 10 cm plastic cell culture dish were transferred to wells using a 1:2 – 1:3 dilution. Cells were further incubated overnight at 37°C and 5% CO_2_ in a cell culture incubator. Before vitrification, grids were removed from the 6-well plate, washed by continuously dripping a total of 10 µl of 10 nm BSA Gold Tracer (Electron Microscopy Sciences) solution and removing the drops with filter paper. Next, 3 µl 10 nm BSA Gold Tracer were pipetted onto the sample side. Using a Vitrobot Mark IV (FEI, now Thermo Fisher), samples were blotted either from both sides or from the backside of the grid by replacing the filter paper on the sample side with a same-sized Teflon sheet, and vitrified by plunging into liquid ethane. See (*22*) for a step-by-step protocol. For cryo-correlative light and electron microscopy imaging, SK-MEL-2 cells expressing CLTA-eGFP were serum starved in medium without fetal bovine serum for 10 – 30 min. Grids were then incubated in a 3 µl drop of medium containing 25 µg/ml Alexa Fluor^®^ 647 ChromPure human transferrin (Jackson Immuno Research Inc.) for 2 – 6 min prior to initiating the washing steps in BSA Gold Tracer solution. Chamber conditions for the Vitrobot Mark IV were set to 37°C and 90% humidity. To ensure that suitable grids were produced for each imaging session, a range of plot forces between 0 and 10, blotting times of 2.5 to 4 s, and a 1 s drain time, were used. Grids were fixed into AutoGrid carrier (Thermo Fisher) and stored in liquid nitrogen until they were imaged.

### Cryo-fluorescence light microscopy

Cryo-samples prepared from single-tagged CLTA-GFP expressing SK-MEL-2 cells and incubated with Alexa Fluor^®^ 647 ChromPure human transferrin (Jackson Immuno Research Inc.) were imaged on a Leica EM Cryo CLEM system (Leica). The system consists of a Leica DM6 FS widefield microscope that is equipped with a motorized Leica EM Cryo stage and a short working distance (<0.28 mm) 50x Leica EM Cryo CLEM ceramic tipped objective (numerical aperture = 0.90). These specifications allow sample imaging at liquid nitrogen temperatures. A halogen lamp powered by a CTR6 halogen supply unit was used as light source. We used GFP ET (Excitation: 470/40, Dichroic: 495, Emission 525/50), Y5 (Excitation: 620/60, Dichroic: 660, Emission 700/75) filter cubes for imaging. For cryo-correlative light and electron microscopy, a grid overview map was recorded using transmitted light and GFP channels. The map was used to identify sufficiently spread cells in regions with good ice quality. The CLTA-GFP signal appeared diffuse in regions of thick or crystalline ice. Z-stacks (total Z = ∼6 µm in 0.6 µm steps) of cells with clear CLTA-GFP foci were recorded using the transmitted light, GFP (CLTA) and Y5 (transferrin) channels. The same imaging conditions were used for cells on individual grids. Imaging conditions were varied between grids to obtain sufficient signal to allow correlation in later steps. Note that some cells or CME sites did not show a transferrin signal most likely either because not all CME sites were loaded with transferrin cargo or due to inconsistent labeling. Images for panel generation in Adobe Illustrator were prepared using FIJI and Adobe Photoshop. No nonlinear gamma correction was applied during image processing.

### Cryo-electron tomography data acquisition

Samples were imaged on a Titan Krios transmission electron microscope (FEI) operated equipped with a X-FEG electron source, a BioQuantum energy filter (Gatan) and a K2 or K3 direct electron detecting device (Gatan) (see supplemental table 1 for details on the used detector) and operated at 300 kV. Samples were visually inspected for ice quality using the FEI flu cam. Overview grid maps were acquired at ∼0.2 µm pixel size. If samples were imaged by cryo-fluorescence light microscopy before, these grid maps were used to identify cells of interest from the cryo-fluorescence light microscopy overview maps. For random data collection, cells with thin cell regions were located. For both types of data collection, polygon maps of the regions of interest with an overlap of 20-25% between individual images were recorded. The polygon maps were used to pick tilt series acquisition points either at random or based on fluorescent signal from cryo-fluorescence light microscopy. The hole pattern of the carbon film was used as a guide. Acquisition points were chosen with adequate distance between individual points to prevent electron dose exposure damage prior to data collection. Only cellular regions without obvious sample damage (e. g., through blotting) were used for data collection. SerialEM (Mastronarde, 2005) in low-dose mode was used for automatic tilt-series recording using a bidirectional tilt scheme starting at +20° and with a typical tilt range from +60° to -60° and base increment of 2°. Pixel sizes between 2.97 and 3.72 Å were used. Target defocus was varied between -2 and -8 µm and the targeted total electron dose was 100 e^-^/Å^2^ (see supplemental table 1 for acquisition parameter details of individual tomograms presented in here). Data was collected in super resolution mode on the K2 detector or in 0.5 binning mode on the K3 detector. Frame time was 0.2 – 0.25 s. Note that at the beginning of the project, the Leica cryo-CLEM microscope was not available to us and we therefore used the double-tagged fluorescent SK-MEL-2 cell line to verify that CME occurs in cells growing on holey carbon grids by fluorescence microscopy.

### Tomogram reconstruction, segmentation model generation and subtomogram averaging

Tilt series alignment and tomogram reconstruction were done using the freely available IMOD software package (Kremer et al., 1996). Tilt series were aligned using 10 nm gold particles as fiducials and tomograms were reconstructed using the backprojection algorithm combined with the SIRT-like filter. Tomograms were then filtered using the Nonlinear Anisotropic Diffusion filter and binned by a factor of 2 using the *binvol* function to further increase contrast. Tomograms from a total of 7 independent datasets were analyzed (see supplemental table 1 for details).

Subtomogram averaging was performed using PEET. The tomograms used are indicated in supplemental table 1 (Heumann et al., 2011; Nicastro et al., 2006). Subtomograms were picked manually and initial motive lists with starting orientations for alignment were generated using the *spikeInit* or the *slicer2MOTL* functions. A single subtomogram was used as the starting reference. Subtomograms were iteratively aligned and averaged and an updated coarse aligned average was used after each iteration. For the subtomogram average of the clathrin hub, 3-fold rotational symmetry was applied. Fourier shell correlation was performed by using the *calcFSC* function and plotted using the *plotFSC* function. ChimeraX (https://www.rbvi.ucsf.edu/chimerax/) was used for subtomogram average visualization and docking of PDB structures. The clathrin hub isosurface model that was used in the segmentation models was generated in IMOD.

Segmentation model generation in IMOD was done by manually tracing actin filaments and membrane shapes and plotting the clathrin hub isosurface model to the original subtomogram positions that were refined during the subtomogram averaging procedure. Branch junctions were identified manually and modeled as a scattered point object (spherical object) that was placed in the center of the junction density.

Tomograms and corresponding segmentation models were cropped to a volume that was centered on the CME site or CCV and had a size of about 500 nm x 500 nm in x and y dimension while the z dimension was kept at its original value. Actin filaments were only considered for further analysis when they were contained in the cropped volume and thus in close proximity to the clathrin-coated feature. For filament length analysis (see below), the original length of the uncropped filament was used if only a part of that filament was contained in the cropped volume.

Images for panel preparation in Adobe Illustrator were generated in IMOD and further processed in Adobe Photoshop. No nonlinear gamma correction was applied during image processing.

### Size measurement of clathrin-coated features

CCV and CCP membrane and coat diameters were measured in the slicer view mode in IMOD. Rotation angles were adjusted to bring the CCV/CCP into full view as a symmetric object. A total of 3 measurements per CCV/CCP were performed on the central tomographic slice and averaged to compensate for imprecision. Measurements for CCPs with a constricted neck were performed at the widest point of the invagination. Further analysis and graph generation were done in Prism8.

### Filament length analysis

Filament length was extracted from the segmentation models using the *imodinfo* function in IMOD. Further analysis and graph generation were done in Prism 8.

### Branch angle measurements

Subtomograms of the branch junctions were extracted using the junction model points and the *boxstartend* function in IMOD. Subtomograms were displayed in the slicer view mode in IMOD. Rotational angles were adjusted to bring both filaments at the junction into view as shown in Fig. 3A and fig. S3A. Tiff images of these views were saved and opened in FIJI. Branch angle measurement was performed using the *Angle tool*. To minimize the effect of filament flexibility, measurements were performed close to the junction and minimal parts of the filaments were included. To assess the effect of the missing wedge, which is most prominent in the z direction, on the branch angle measurements, we plotted the measured branch angle against the x and y rotations that had to be applied to get the junction in view. The equal distribution of branch angles across the rotation angles shows that our measurements were not affected by the missing wedge at this level of accuracy.

### Filament polarity and orientation analysis

Branched actin filament polarity was analyzed based on branch junction geometry as indicated in Fig. 3A. For analysis of unbranched filament polarity, we adapted a previously published cross-correlation based method (Narita et al., 2012). For the cross-correlation analysis, we first generated an artificial 3D actin filament from a previously published structure (PDB ID: 6DJM; (Chou and Pollard, 2019)) using ChimeraX and IMOD. We then used the *relion_image_handler* function in Relion to rescale and lowpass filter the artificial filament to the pixel size of our binned tomography data (5.943 Å) and a resolution of 30 Å. Next, a tiff image stack of 2D projection images of the artificial actin filament was generated using the *xyzproj* function in IMOD. Each of the projection images was taken after rotating the filament by 5° around the filament axis (Φ) relative to the previous image. The complete image stack contained a total of 36 images covering a 180° rotation of the artificial actin filament. The image stack was then cropped to 69 pixels x 23 pixels, corresponding to the length of a 13-subunit actin filament. This size was picked because of the structural organization of actin filaments. The symmetry of an actin filament can be described as a single strand left-handed helix with about 13 monomers repeating every six turns (Dominguez and Holmes, 2011). Accordingly, a 13-subunit reference projection series contains sufficient information about the helical organization of actin filaments to serve as a reference structure. We then generated a second reference image stack with opposing polarity by rotating the original stack by 180° (fig. S4A). Hereafter we refer to the reference with the plus pointing up as ref+_angle-i_ and the one with the minus end pointing up as ref-_angle-i_. Filaments for polarity analysis were extracted from the tomograms using the corresponding segmentation models and the *imodmop* function in IMOD. 2D projection images of these filaments were generated in FIJI. Images were oriented as shown as in fig. 4SB and cropped so the resulting image was 23 pixels wide (= width of the reference images). Note was taken on the rotation that was applied to the images, and only straight parts of the filaments were included. From these test filaments (test), sub-images of the size of the reference images (69 pixels x 23 pixels) were extracted with a frequency of 5 pixels (test_i_), corresponding to about one actin subunit in the filament (fig. S4B). The cross-correlation coefficient (R) was calculated for each of the test sub-images test_i_ with each of images in both of the reference stacks ref+_angle-i_ and ref-_angle-i_ (fig. S4C). Then, the correlation curves with ref+_angle-i_ and ref-_angle-i_ were subtracted from each other and a difference curve was generated (fig. S4D). The mean average difference was calculated from these values. If the value was negative, the tested filament was determined to have the same polarity as ref-_angle-i_, if the value was positive, the tested filament was determined to have the same polarity as ref+_angle-i_. The cross-correlation calculation algorithm was prototyped on ImageJ with the Image CorellationJ plugin (https://www.gcsca.net/IJ/ImageCorrelationJ.html). To increase throughput and ease of use, the software was automated and reimplemented into a custom python-based command line tool. The accuracy of this method depends on test filament length. Therefore, only unbranched filaments of at least 80 nm length (> two 13-subunit repeats) were included in the analysis. Branched filaments were included without length consideration because the filament junction identifies the polarity.

To calculate and display the orientation of the filaments in the tomograms based on the determined polarity (either by branch junction geometry or cross-correlation analysis), we developed an analysis pipeline with Jupyter notebooks (Python 3.7) relative to a user-defined reference plane. In general, the reference plane was parallel to the plasma membrane for CME invaginations or tangential to the surface for CCVs. The reference plane was manually segmented using IMOD. We calculated a vector normal to the reference plane, such that 0° orientation was defined as the vector pointing from the filaments toward the reference plane, and 180° was identified as pointing away from the plane. Filaments with model points crossing within 10 pixels (6 nm) from the reference plane were excluded from the analysis. The directionality of the filaments (from minus end to plus end) was calculated as the arccosine of the dot product between the filament vector and the vector normal to the reference plane. Filament positions and orientations were plotted using matplotlib (3.0.2). Further analysis and graph generation were done in Prism8.

### Mathematical modeling using Cytosim

We used the agent-based model in Akamatsu et al., 2020 to run 3D stochastic simulations of the mammalian endocytic actin network (*8*). We used similar parameters and initial conditions with the following modifications:

To test effects of the branch angle variability on force production and internalization efficiency, we changed the average branching angle of the Arp2/3 complex to be 70° rather than 77°, to closer match the measured branch angle in the tomograms (*8*). In both cases the branch flexibility was set to 0.076 pN µm/rad. In general, to average multiple stochastic simulations, we ran 24 simulations per condition. To add Hip1R molecules at the neck, we repurposed Matlab code from Akamatsu et al. 2020 in order to distribute Hip1R molecules uniformly around a cylinder of radius 30 nm to match the shape and diameter of a typical CME neck. Sixty Hip1R molecules were distributed along 45 nm of the surface of the neck. This conservative value for the number of Hip1R molecules at the neck corresponds to a lower molecular surface density relative to the bud, and results in more tractable simulation runs that proceed to completion.

## Acknowledgments

We would like to thank Elizabeth Montabana, Daniel Toso and Jonathan Remis for technical assistance with cryo-ET method development and data collection, Paul Tobias for computation support and Sun Hae Hong for CLTA-GFP cell line generation and validation. Cryo-ET sample preparation was optimized at the LBNL cryo-EM resource and image data was collected at the Bay Area Cryo-EM facility at the University of California, Berkeley. Daniel Fletcher provided valuable remarks on the manuscript. RV thanks the Allen Institute for Cell Science founder, Paul. G. Allen, for his vision, encouragement and support.

## Author contributions

Conceptualization: DS, MA, PR, DGD; Methodology: DS, MA; Investigation: DS, MA, AM, KV, JH; Visualization: DS; Funding acquisition: DS, MA, KMD, PR, DGD; Project administration: DS, DGD; Software: DS, RV, MA, JS; Supervision: DGD, PR; Writing – original draft: DS; Writing – review & editing: DS, MA, PR, DGD.

## Data and materials availability

All data are available in the main text or the supplementary materials. Raw data and cell lines will be made available upon request. Code is deposited on GitHub and will be made available upon request.

