## Supplemental Figures, Table and movie captions for "Actin force generation in vesicle formation: mechanistic insights from cryo-electron tomography"

**This document includes:**

Figure Legends S1 to S5
Table S1
Legends for Movies S1 to S5

**Other Supplementary Materials for this manuscript include the following:**

Movies S1 to S5

### Supplementary Figures

#### Supplementary Figure 1

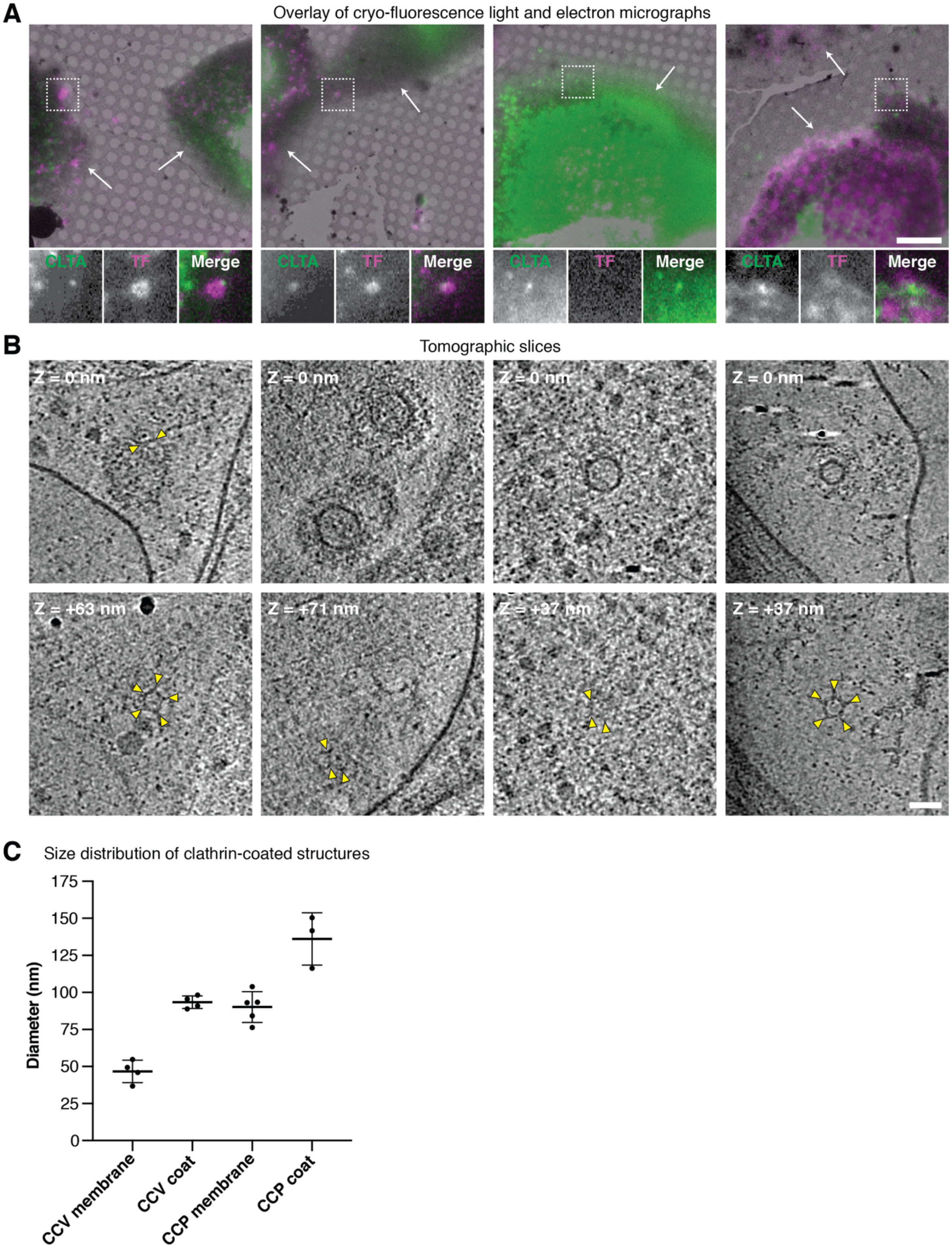

**Fig. S1. Identification of clathrin coats using cryo-CLEM.** (**A**) Overlay of correlative cryo-FLM and cryo-TEM images showing SK-MEL-2 cells endogenously expressing CLTA-GFP (white arrows) growing on EM grids. Fluorescent transferrin (TF, magenta) was added to the cells 2-6 min before freezing. Insets show 2.3x magnified views of boxed regions. (**B**) Slices of tomograms recorded in the boxed regions in (**A**). The EM images are arranged in the same order as the boxed regions in (**A**). Yellow arrowheads point at clathrin coats. (**C**) Diameter distribution of the membrane compartment and clathrin coat of CCVs and CCPs. Thick lines represent mean values, error bars show the standard deviation (SD). Scale bars, 10 µm in (**A**), 50 nm in (**B**).

**Supplementary Figure 2**

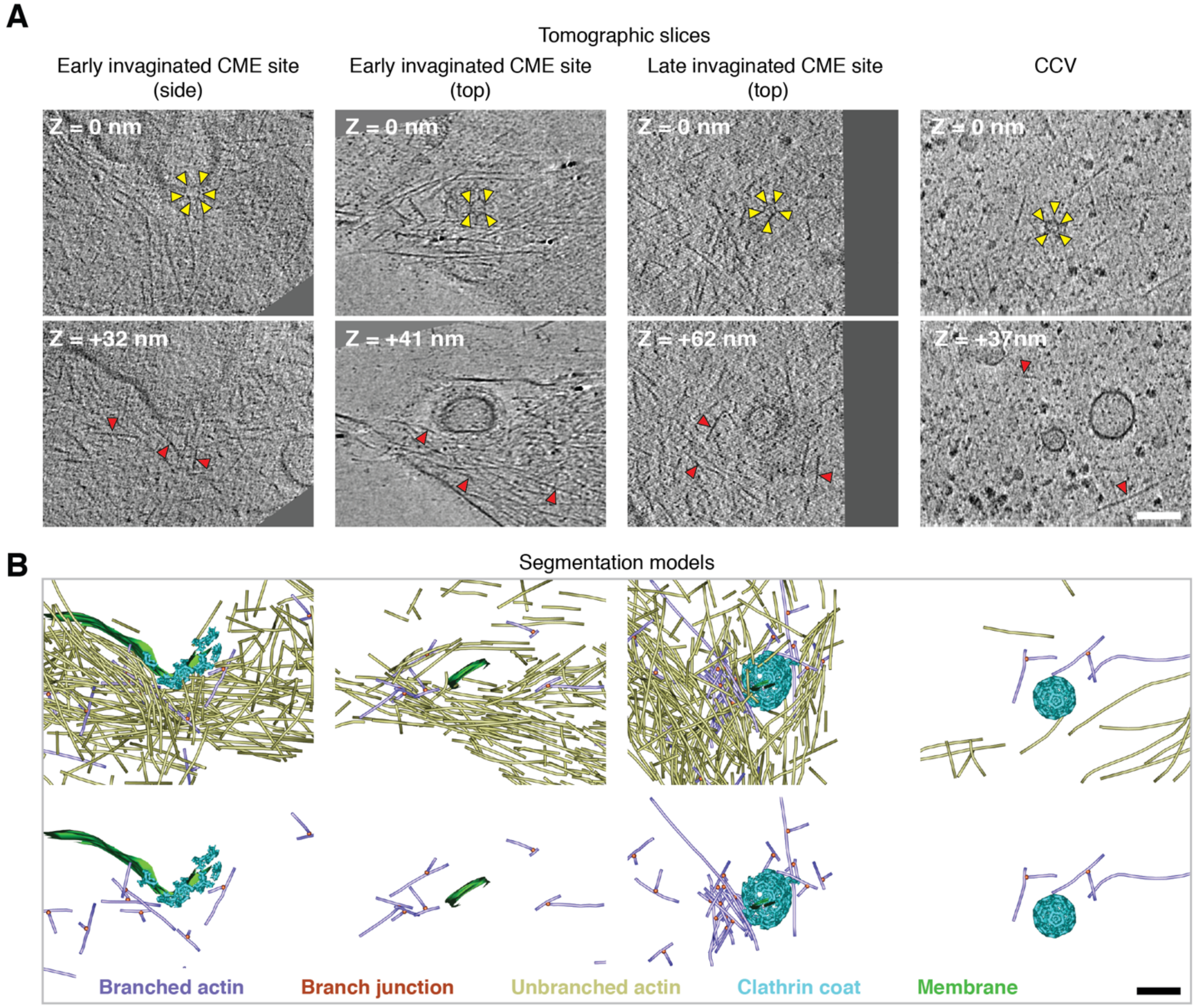


**Fig. S2. Additional examples for branched and unbranched actin filament organization at CME sites.** (**A, B**) Tomographic slices and related segmentation models of the tomograms. Color coded legend describes elements shown in the models. Column 1 shows CME site with orthogonal filament organization relative to the plasma membrane, column 2 and 3 show parallel filament organization. Column 4 shows a CCV. Tomogram in column 1 without the corresponding segmentation model and further analysis shown here was used before in Akamatsu et al., 2020 (*8*). Scale bars, 50 nm in (**A, B**).

**Supplementary Figure 3**

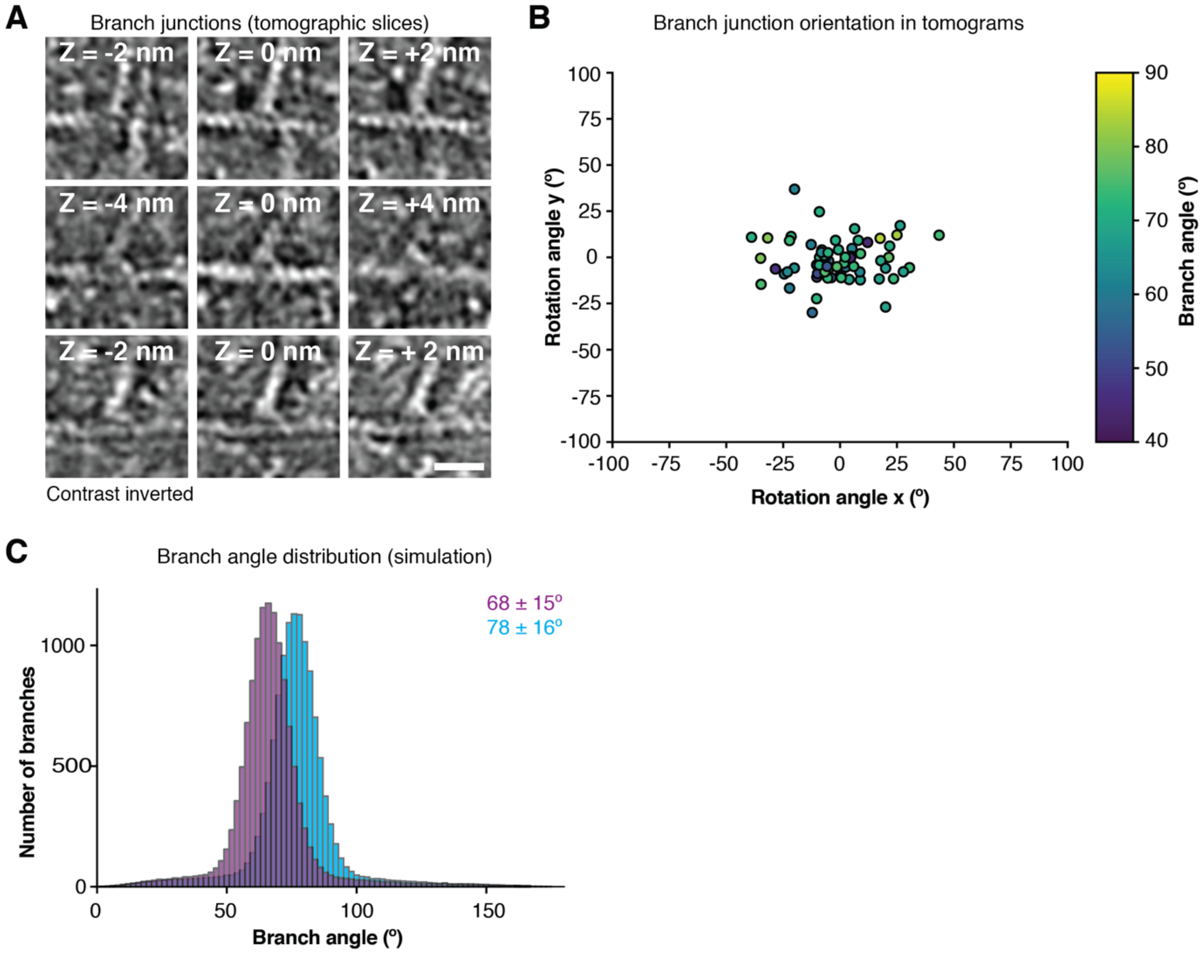


**Fig. S3. Branch junction appearance and orientation in tomograms and branch angle distribution in mathematical model.** (**A**) Tomographic slices of 3 individual branch junctions at Z-positions, left to right, relative to central slice (Z = 0 nm). (**B**) Branch angle measurement dependence on the orientation of the branch junction in the tomogram. Rotation angles describe how much tomographic data had to be rotated to generate the view displayed in (**A**). (**C**) Branch angle distribution obtained from the simulations shown in Fig. 3 (**C**). The initial value of the parameters used were 70º or 77º. In both cases the branch flexibility was set to 0.076 pN µm/rad. Scale bar, 10 nm in (**A**).

Supplementary Figure 4

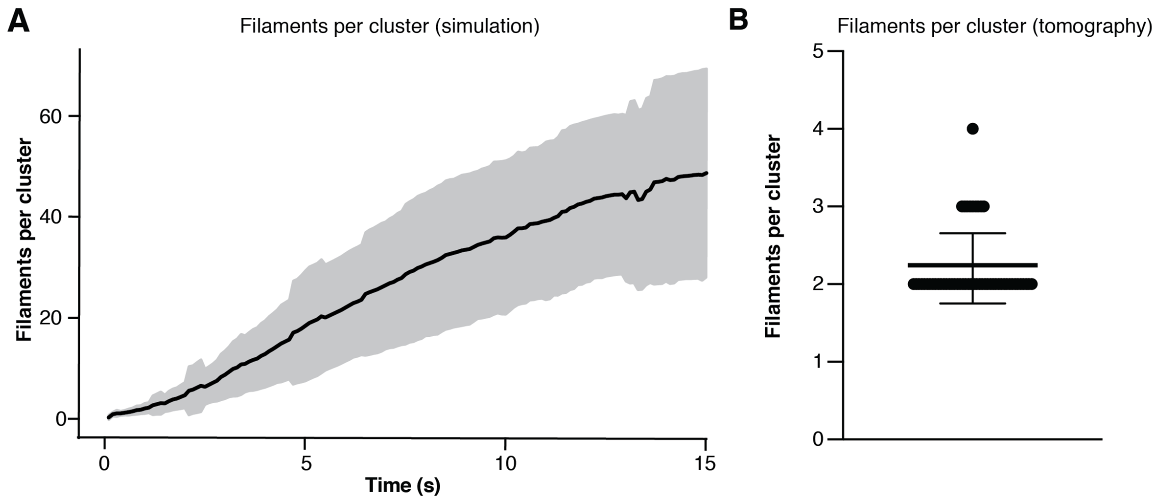


**Fig. S4. Branched actin organization in cryo-electron tomograms and mathematical model.** (**A**) Simulated number of filaments per cluster in the mathematical model. Graph shows mean and SD. (**B**) Number of filaments per cluster in tomograms. Mean and SD are shown.

**Supplementary Figure 5**

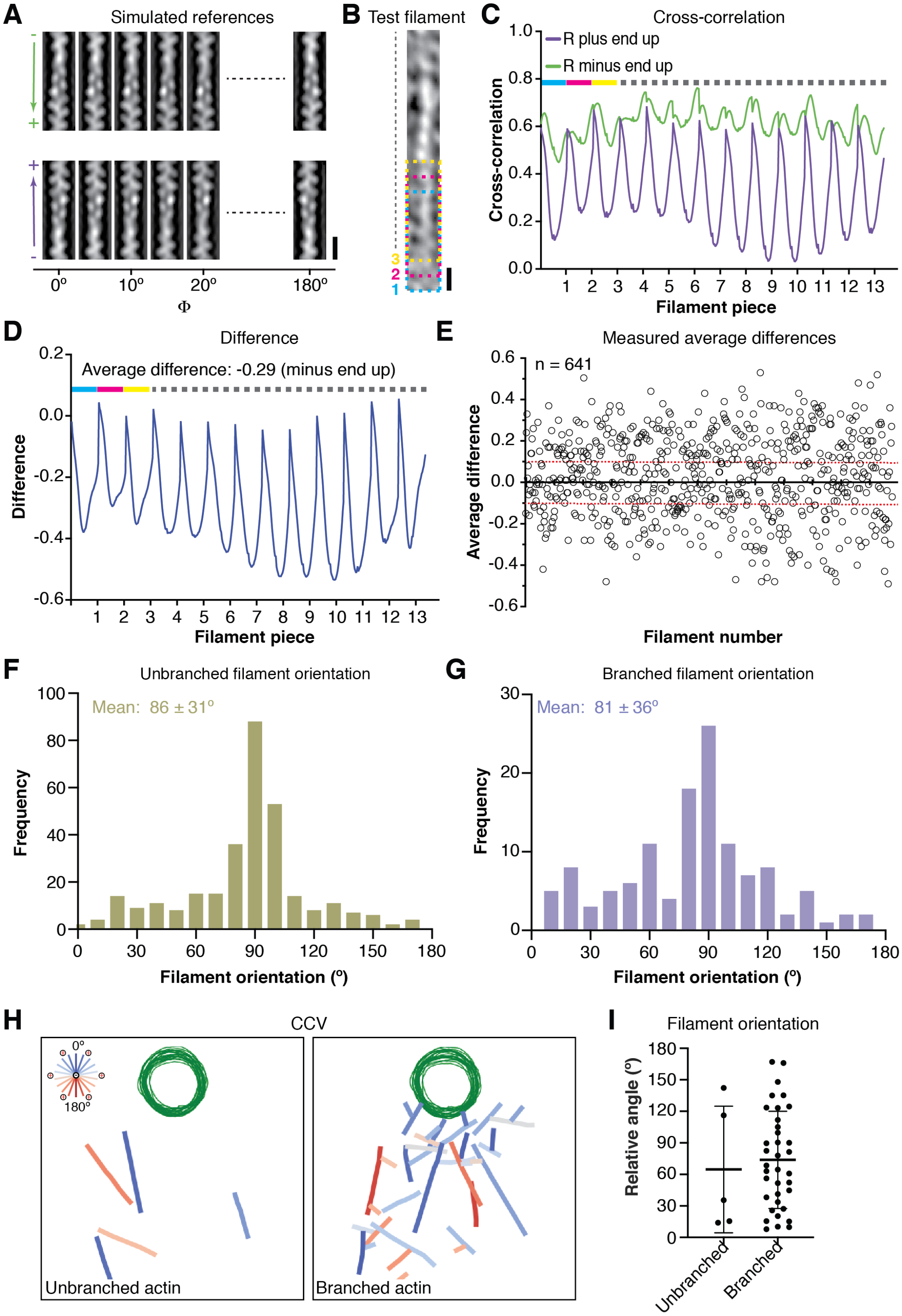

**Fig. S5. Actin filament orientation analysis.** (**A**) – (**E**) Procedure for orientation analysis based on cross-correlation analysis. (**A**) Two sets of reference projection images from artificial actin filaments with opposing polarity at 30 Å resolution. Individual images where obtained after rotating filaments along the filament axis (Φ) by 5º. (**B**) Example filament extracted from a tomogram. Multiple consecutive sections of the filament that differ approximately by the height of one actin subunit are cropped to have the same pixel size as the reference images. Example segments are highlighted by the three colored boxes (**C**) Cross-correlation readout of the filament in (**B**) against the reference. Each curve represents cross-correlation of all pieces of the filament (**B**) against one of the references. Color-coded legend corresponds to the respective color-coded segment in (**B**). (**D**) Difference plot of the curves in (**C**). The average difference indicates the minus end of the filament in (**B**) is pointing upwards. (**E**) Average difference for filaments across all analyzed tomograms. Note that only filaments of at least 80 nm and with an average difference lower than -0.1 or higher than 0.1 were considered for further analysis. (**F**, **G**) Unbranched (**F**) and branched (**G**) actin filament orientation distribution at CME sites across all tomograms relative to the normal (0º) of a reference plane representing the position of the plasma membrane. (**H**) Color-coded filament orientation in proximity to a CCV (green). (**I**) Relative filament to reference plane orientation to CCV shown in (**H**). Mean and SD are shown.

Scale bars, 10 nm in (**A**, **B**).

Supplementary Figure 6

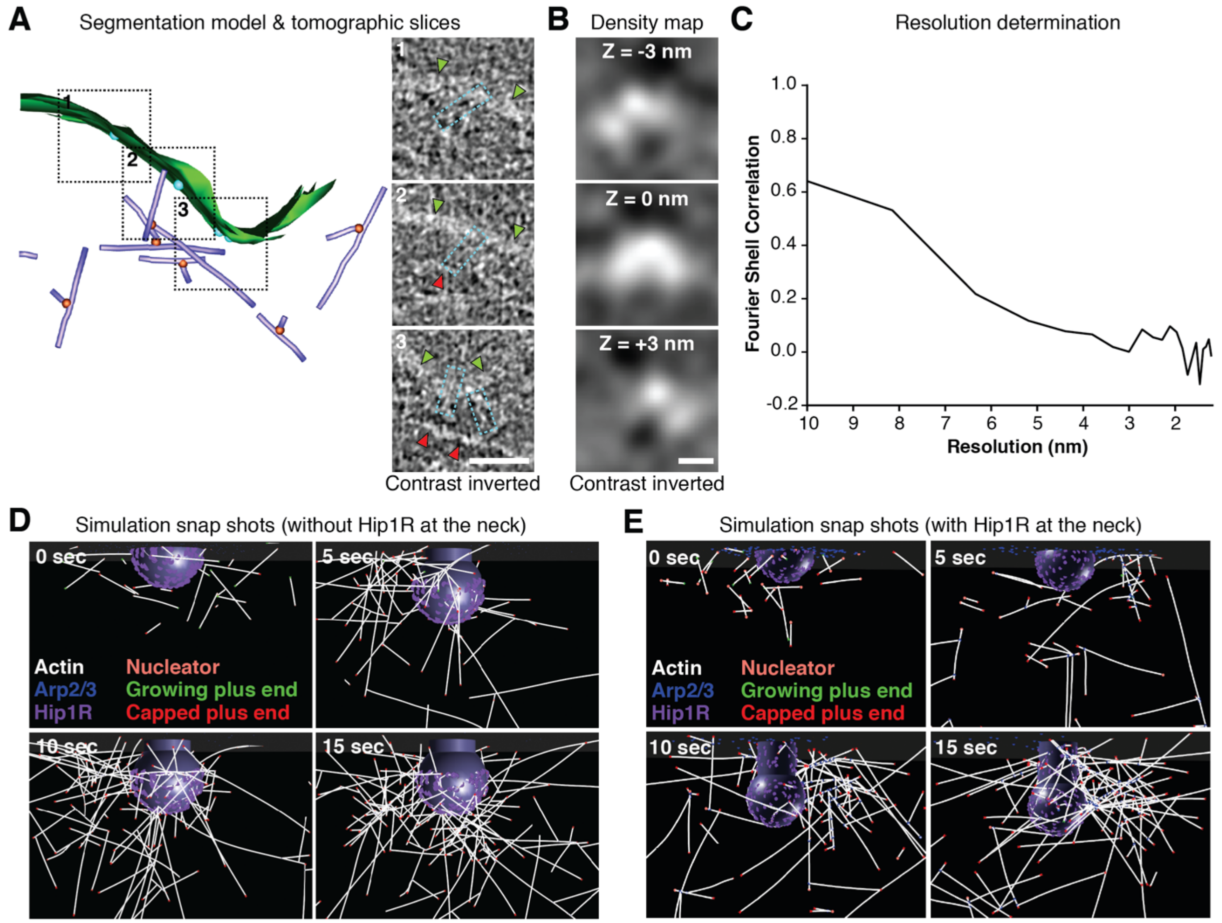


**Fig. S6. Identification and distribution of putative Hip1R dimers at CME sites.** (**A**) Tomographic slices of the regions indicated in the segmentation model of the tomogram shown in fig. S2A, B. Slices highlight putative Hip1R dimers (white boxes) that are associated with the coated membrane of the CME invagination (green arrowheads) and their positions are highlighted by cyan spheres in the segmentation model correspond. Red arrowheads point at actin filaments associated with a putative Hip1R dimer. (**B**) Density map of the putative cytoplasmic actin binding domain of a Hip1R dimer obtained by subtomogram averaging. (**C**) Resolution determination of the density map in (**B**) based on FSC. (**D**. (**E**) Snapshots of model simulation (**D**) without and (**E**) with Hip1R at the neck region. Note enrichment of actin filaments at the neck of the invagination. Scale bars, 50 nm in (**A**), 5 nm in (**B**).

#### Supplementary Table

##### Table S1. Analyzed tomograms and data collection details

| Data set | CME stage | Detector | Pixel size (Å) | Target defocus (µm) | Correlative data (y/n) | Used for clathrin subtomogram averaging (y/n) |
|---|---|---|---|---|---|---|
| 1 | Late invagination* | K2 | 3.72 | -8 | n | n |
| 2 | CCV* | K2 | 2.97 | -6 | n | y |
| 2 | CCV | K2 | 2.97 | -6 | n | y |
| 3 | Late invagination* | K2 | 2.97 | -2 | n | y |
| 3 | Early and late invagination* | K2 | 2.97 | -2 | n | y |
| 3 | Late invagination | K2 | 2.97 | -2 | n | y |
| 4 | Early/flat site* | K2 | 2.97 | -6 | n | y |
| 5 | CCV* | K2 | 2.97 | -2 | n | y |
| 6 | Early invagination | K3 | 3.07 | -6 | y | n |
| 6 | Undetermined (likely late invagination or CCV) | K3 | 3.07 | -6 | y | n |
| 6 | Undetermined (likely late invagination or CCV) | K3 | 3.07 | -6 | y | n |
| 7 | CCV | K3 | 3.07 | -6 | y | n |

*Segmented tomograms presented in Fig. 2 and fig. S2

**Supplementary Movie Legends**

**Movie S1.**

Density map and surface rendering model of clathrin hub subtomogram average. Scale bar, 5 nm. Frame rate, 15 frames per second.

**Movie S2.**

Cryo-electron tomogram and segmentation model of early CME site covered by dense cortical actin filament network. Scale bar, 50 nm. Frame rate, 10 frames per second.

**Movie S3.**

Cryo-electron tomogram and segmentation model of late invaginated CME site with orthogonal branched actin filament organization relative to the plasma membrane. Scale bar, 50 nm. Frame rate 10 frames per second.

**Movie S4.**

Cryo-electron tomogram and segmentation model of late invaginated CME site with parallel branched actin filament organization relative to the plasma membrane. Note dense actin cortex around CME site. Scale bar, 50 nm. Frame rate 10 frames per second.

**Movie S5.**

Cryo-electron tomogram and segmentation model of clathrin-coated vesicle and associated actin tail. Scale bar, 50 nm. Frame rate 5 frames per second.
